# Carcinogenicity and testicular toxicity of 2-bromopropane in a 26-week inhalation study using the rasH2 mouse model

**DOI:** 10.1101/2022.11.06.515381

**Authors:** Yuko Goto, Arata Saito, Kenji Takanobu, Hideki Senoh, Misae Saito, Yumi Umeda, Shotaro Yamano

## Abstract

2-Bromopropane (2-BP) is a colorless liquid at room temperature and is used in closed systems in factories, mainly as an intermediate for medicines, pesticides, and other chemicals. However, the carcinogenicity of 2-BP is still unknown. The CByB6F1-Tg(HRAS)2Jic (rasH2) transgenic mouse model has been established as an alternative to long-term studies (1.5years-lifetime) to detect carcinogenicity in as short a time as six months. We performed a 26-week inhalation exposure study of 2-BP using the rasH2 mouse model. Male and female rasH2 mice were exposed to 0, 67, 200, or 600 ppm of 2-BP for 6 hours/day, 5 days/week for 26 weeks. All tissues and blood were collected and subjected to biological and histopathological analyses. The results showed a concentration-dependent increase in lung tumor development in male and female rasH2 mice exposed by inhalation to 2-BP, which was significant by Peto’s trend test. Furthermore, in male rasH2 mice, 2-BP was found to be a testicular toxin. This study is the first to demonstrate that 2-BP is carcinogenic in male and female mice and a testicular toxin in male mice in the rasH2 mouse model.

## Introduction

2-Bromopropane (CAS number: 75-26-3, hereafter 2-BP) is a colorless liquid at room temperature and is used in closed systems in factories, mainly as an intermediate for medicines, pesticides and other chemicals. The Montréal Protocol was adopted in Canada in 1987 to regulate the production, consumption, and trade of ozone-depleting chemicals such as Chlorofluorocarbons (CFCs) ^1^. In addition to chemical manufacturing, since the adoption of the Montréal Protocol the use of 2-BP as a solvent for cleaning has increased in various factories as an alternative to CFCs and 1,1,1-trichloroethane. In the fall of 1995, an outbreak of menstrual arrest and pancytopenia among female workers and oligozoospermia or azoospermia among male workers exposed to a 2-BP cleaning solution at an electronic parts factory in South Korea was reported ^2,3^. The causative agent was 2-BP, whose toxicity to the testes, ovaries, and hematopoietic organs has been reproduced in numerous experiments using rodents ^4–18^. However, there is little information on the carcinogenicity of 2-BP. Studies examining carcinogenicity in test animals have not been reported. *In vitro* studies have reported that 2-BP induces mutagenicity in Salmonella typhimurium TA100 and TA1535, but mutagenicity using Escherichia coli WP2 uvrA and chromosome aberration using Chinese hamster lung cells were both negative ^19^.

Carcinogenicity studies on food additives, existing carcinogens, and oral drug candidates have demonstrated that rasH2 mice are more sensitive to genotoxic as well as non-genotoxic carcinogens than a p53 heterozygous mouse model^20–23^. Consequently, the CByB6F1-Tg(HRAS)2Jic (rasH2) transgenic mouse model can be used for detecting carcinogenicity in as short a time as six months. This model has been established as an alternative to long-term studies (1.5 years-lifetime) to predict the carcinogenic potential of chemicals^24^.

We have previously conducted systemic inhalation exposure studies of various chemicals in rodents, including rasH2 mice, and have examined various toxicities, including carcinogenicity and pulmonary fibrosis ^25–27^. In this study, we conducted a 26-week systemic inhalation exposure study of 2-BP using rasH2 mice and comprehensively evaluated carcinogenicity in each organ and reproductive organ toxicity.

## Material and Methods

### Ethics declarations

All animals were maintained and used in accordance with Guidelines for the Care and Use of the Institutional Animal Care and Use Committee of the Japan Bioassay Research Center. All the animal experiments were approved by the Institutional Animal Care and Use Committee (Approval No.: 0164) and Genetic Recombination Experiments Safety Committee (Approval No.: 2016-02) of the Japan Bioassay Research Center, and have been reported following the recommendations in the ARRIVE guidelines. We have complied with all relevant ethical regulations for animal testing and research.

### Materials

2-Bromopropane (2-BP) 1^st^ grade (Lot No.: TWQ6860) was purchased from Wako Pure Chemical (Osaka, Japan). Their detailed characteristics of 2-BP 1^st^ grade is given in Fig. 1A. 2-BP was analyzed by mass spectrometry equipped with direct probe (M-80B, Hitachi, Tokyo, Japan) before its use and analyzed by gas chromatography (5890A, Agilent Technologies, Santa Clara, CA) before and after its use to confirm its stability and purity. Gas chromatography was performed as follows: column: G-950 (1.2 mmφ, 20 m); column temperature: 150□; Flow rate: 10 mL/min; Detector: Flame ionization detector. Other reagents used in the study were of the highest grade available commercially. Additionally, 2-BP in the chambers was monitored using gas chromatography (14-B, Shimadzu, Kyoto, Japan): no gas chromatographic peaks other than 2-bromopropan were detected in the inhalation exposure chambers.

**Fig. 1.**
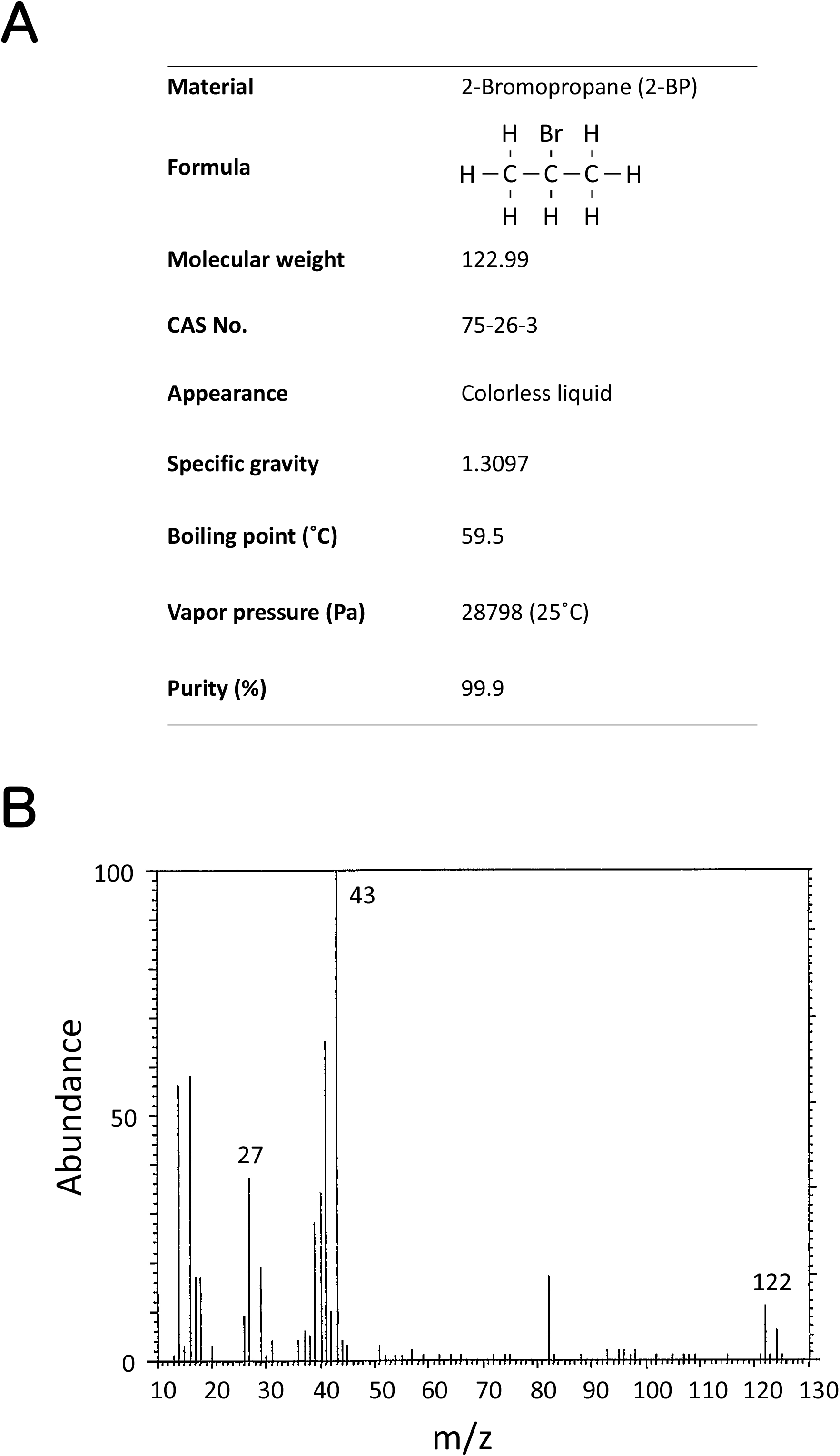
Characterization of 2-bromopropane (2-BP). A: General properties. B: Mass Spectrum.

### Animals

6 weeks old male and female rasH2 mice were purchased from CLEA Japan, Inc. (Tokyo, Japan). Mice were housed in an air-conditioned room under a 12 hour light/12 hour dark (8:00-20:00, light cycle) photoperiod, and fed a Certified Diet (CRF-1, Oriental Yeast Co. Ltd., Tokyo, Japan) and tap water *ad libitum*. After 6 days of quarantine and 5 days of acclimatization, they were exposed to 2-BP from 8 weeks of age by the procedure described below.

### Generation of 2-bromopropane

This study referred to the Organisation for Economic Co-operation and Development (OECD) guidelines for carcinogenicity studies (OECD TG 451) ^28^. Four inhalation exposure chambers (volume: 3.7 m^3^) were used throughout the 26-week exposure period. 2-BP in a reservoir flask with a thermostatic water bath that was controlled at 22□. Clean air was bubbled through liquid 2-BP to generate a saturated vapor mixture. Airflow containing the saturated 2-BP vapor was cooled to 17□ by passing it through a thermostatic condenser, resulting condensation of 2-BP. Then the liquid 2-BP was re-warmed to 2□, vaporizing the 2-BP, and diluted with clean air. Finally, the diluted vapor-air mixture was then introduced into the spiral line mixer of each inhalation exposure chamber, with the flow rate of the vapor-air mixture into each spiral line mixer being adjusted using a flow meter for each target concentration. The spiral line mixer diluted the 2-BP vapor-air mixture to the target concentrations with clean air.

### 26-week inhalation study

This experiment was conducted with reference to “Standards to be Observed by Testing Institutions” Notification No. 76 of the Ministry of Labour, Japan, 1 September 1988 (amendment: Notification No. 13 of the Ministry of Labour, Japan, 29 March 2000) and reference to the OECD principle of Good Laboratory Practice ^29^. For dosage setting, we conducted a preliminary study of 4-weeks inhalation exposure to 2-BP (100, 300, 1000, and 3000 ppm) using the non-TG rasH2 mice in accordance with the OECD TG 412 ^30^. A suppression in weight gain was observed among the males and females in the 1000 and 3000 ppm exposure group. Therefore, the maximum target concentration of 2-BP in the 26 week study was set at 600 ppm (twice the no-effect level of the 4 week study): target concentrations for 2-BP were set at 67, 200 and 600 ppm. The exposure schedule was 6 hours per day; 5 days per week, for 26 weeks. Two hundred mice with 25 males and 25 females in each group were housed in individual stainless-steel cages and maintained at a temperature of 23 ± 2°C and a relative humidity of 50 ± 20% and a 12-h light/dark cycle with 12 air changes per hour during the non-exposure periods and 7-9 air changes per hour during the exposure periods. During exposure to 2-BP for 6 hours, the mice had free access to food and water. During the study period, body weight and food consumption were measured once a week. All animals were fasted from the day before the autopsy date. At 1-4 days after the last exposure, the blood of the mice was collected under isoflurane anesthesia and the mice were euthanized by exsanguination. Organs including the adrenal, thymus, testis, ovary, heart, lung, spleen, liver, kidney, and brain were weighed, and all organs were examined for macroscopic lesions. For histopathological analysis, all the tissues were collected from all mice in each group and fixed in 10% neutral phosphate buffered formalin solution. The right lung was directly fixed by immersion. The left lung was inflated with fixative at a water pressure of 20–25 cm, and then fixed by immersion.

### Hematological and blood chemistry tests

For hematological examination, blood samples collected at the time of each autopsy were analyzed with an automated hematology analyzer (ADVIA120, Siemens Healthcare Diagnostics Inc. Tarrytown, NY). For biochemical tests, the blood was centrifuged at 3,000 rpm (2,110×*g*) for 20 minutes, and the supernatant was analyzed with an automated analyzer (Hitachi 7080, Hitachi, Ltd., Tokyo, Japan).

### Histopathological analysis

Tissue sections were cut from paraffin-embedded specimens of all organs required by OECD TG451 ^28^, and the section (2-μm thick) was stained with hematoxylin and eosin (HE) for histological examination. The histopathological finding terms used in this study for lesions were determined by certified pathologists from the Japanese Society of Toxicologic Pathology, and based on the finding terms adopted by International Harmonization of Nomenclature and Diagnostic Criteria for Lesions in Rats and Mice (INHAND) ^31^. Pathological diagnosis was performed blindly by three pathologists and summarized after a cumulative discussion.

### Statistical analysis

All statistical analysis was carried out by the BAIS system (Hitachi Social Information Services Ltd., Tokyo, Japan) and GraphPad Prism 5 (GraphPad Software, San Diego, CA). The incidences of non-neoplastic lesions were analyzed using the chi-square test, and the severity was defined as 1, slight; 2, moderate; 3, marked; and 4, severe. The incidence of neoplastic lesions was statistically analyzed by Fisher’s exact test and incidence trend was analyzed by Peto’s trend test. Body weight, organ weight, food consumption, and hematological and blood biochemical parameters were analyzed by Dunnett’s multiple comparison test. All statistical significance was set at *p*<0.05.

## Results

### Characterization and concentration in the inhalation chamber of 2-Bromopropane

The test substance was analyzed by mass spectrometry before its use. The measured mass spectrum (Fig. 1B) was consistent with the literature spectrum of 2-bromopropane (2-BP) ^32^. 2-BP was also analyzed by gas chromatography for purity and stability before and after its use. Gas chromatography indicated one major peak before and after its use. No impurity peaks were detected before or after its use (data not shown), indicating that the 2-BP used in the present study was stable for the duration of the study period.

The 2-BP concentrations were at the target concentrations over the 26-week exposure period: 66.8 ± 1.2 ppm for the 67 ppm group, 200.6 ±3.6 ppm for the 200 ppm group, and 599.2 ± 10.0 ppm for the 600 ppm group.

### Clinical findings, survival, body weight curve, food intakes and final body weight

Observations of the general condition of the mice and clinical findings showed no changes throughout the study period that could be attributed to the effects of 2-BP. There were no significant changes in the survival of females or males exposed to any concentration of 2-BP compared to the control groups (Fig. 2A, B). Suppression of body weight gain by males exposed to 200 ppm and 600 ppm 2-BP and females exposed to 600 ppm was observed (Fig. 2C, D), and there was a statistically significant reduction in final body weight in males exposed to 200 and 600 ppm and females exposed to 600 ppm compared to the control group (Fig. 3A, B). Changes in food intake were not as pronounced as those in body weight, and no concentration-dependent changes were observed in either sex (Fig. 2E, F).

**Fig. 2.**
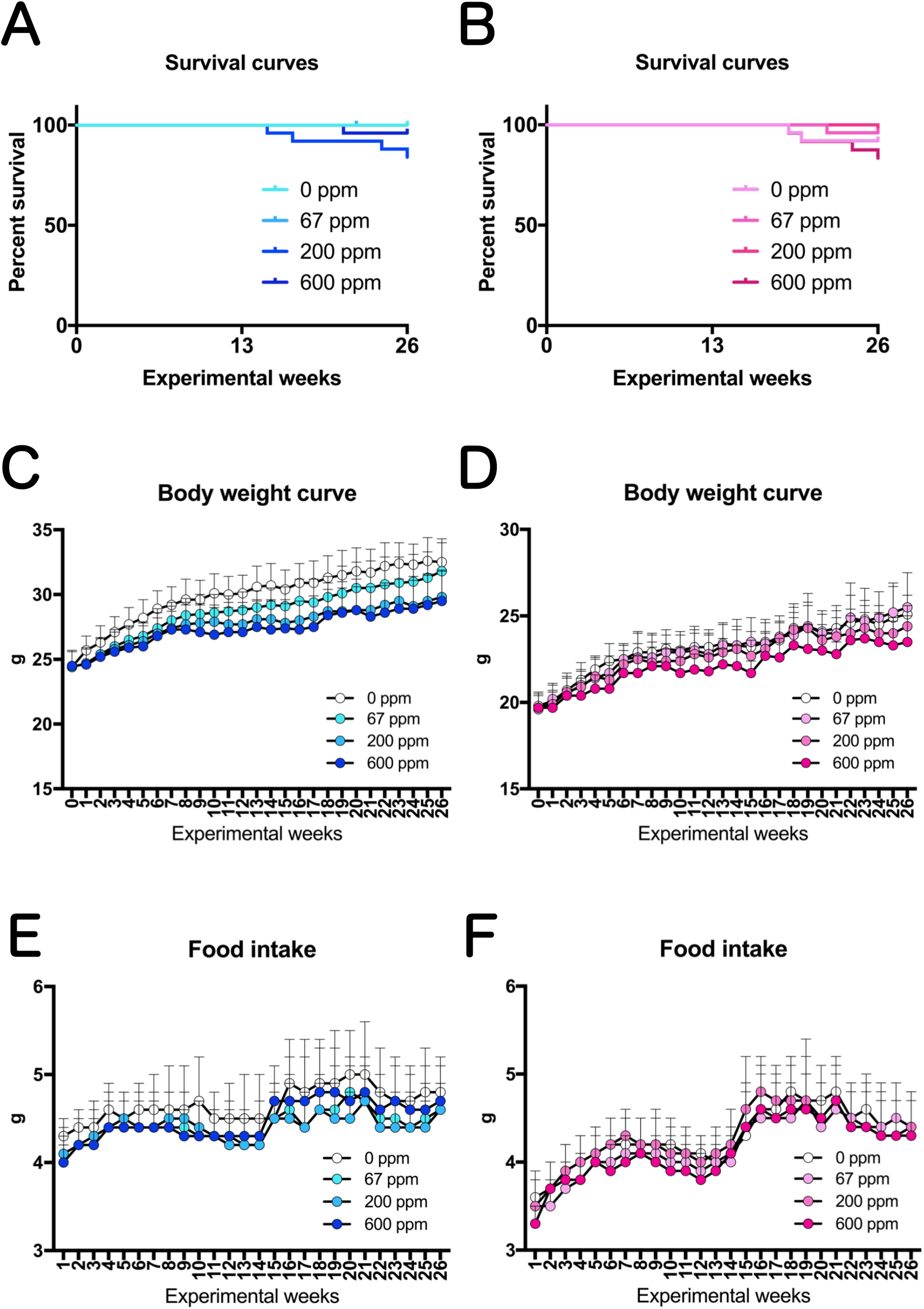
Survival curves, Body Weight curves, and Food Intake of rasH2 mice exposed by inhalation to 2-BP (67, 200, or 600 ppm, 6 hours/day, 5 days/week, 26 weeks). A, C, E: Male mice. B, D, F: Female mice.

**Fig. 3.**
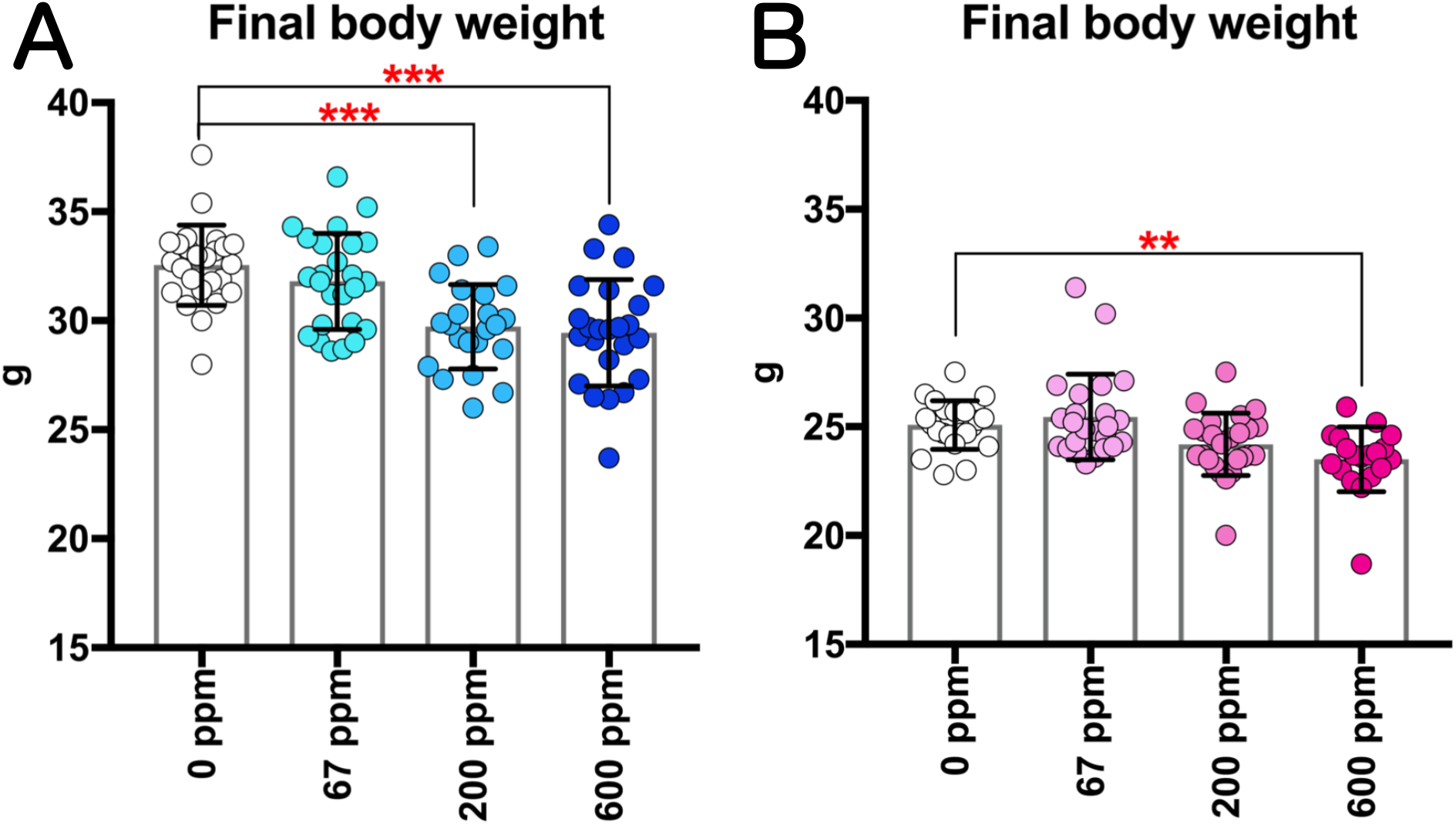
Changes in the final body weights of rasH2 mice following inhalation exposure to 2-BP (67, 200, or 600 ppm, 6 hours/day, 5 days/week, 26 weeks). Final body weights in males (A) and females (B) were measured at sacrifice. Dunnett’s multiple comparison test was used to compare weights with the age-matched control (0 ppm) groups: \*\**p*<0.01 and \*\*\**p*<0.001.

### Carcinogenicity of 2-BP in rasH2 mice

In this study, tumors were observed in several organs of male and female rasH2 mice, including the lung, skin, lymphatic vessel, oral cavity, stomach (forestomach), liver, Harderian gland, urinary bladder, lymph node, thymus, subcutaneous tissue (subcutis), spleen, bone marrow, nasal cavity, and vagina. A summary of the histopathological findings for the neoplastic lesions in all organs is given in Table 1. Both male and female rasH2 mice had numerous bronchiolo-alveolar adenomas and carcinomas in the lungs, one of the predominant tumor organs in rasH2 mice. Representative microscopic photographs of lung tumors are shown in Fig. 4. The adenomas that developed in 2-BP exposed rasH2 mice (Fig. 4A) exhibited typical adenoma histology with the solitary adenoma nodule compressing the surrounding tissue (Fig. 4B) and being composed of cells with large nuclei and high nuclear/cytoplasmic ratios (Fig. 4C) compared to normal alveolar epithelium (Fig. 4D) which constitutes the surrounding alveolar tissue. The carcinoma shown in Figure 4E occupies a single lung lobe (Fig. 4E) and has disseminated into the alveolar air space (Fig. 4F) and bronchial air space (Fig. 4G). The presence of tumor cells in the airspace was also observed in the right lung, which was fixed without formalin injection from the bronchus, and thus is not an artifact of formalin injection. Occasionally, carcinoma tumor cells metastasized to other lung lobes (intrapulmonary metastasis) (Fig. 4H). There were no differences in the histological characteristics of these tumors due to 2-BP exposure.

**Fig. 4.**
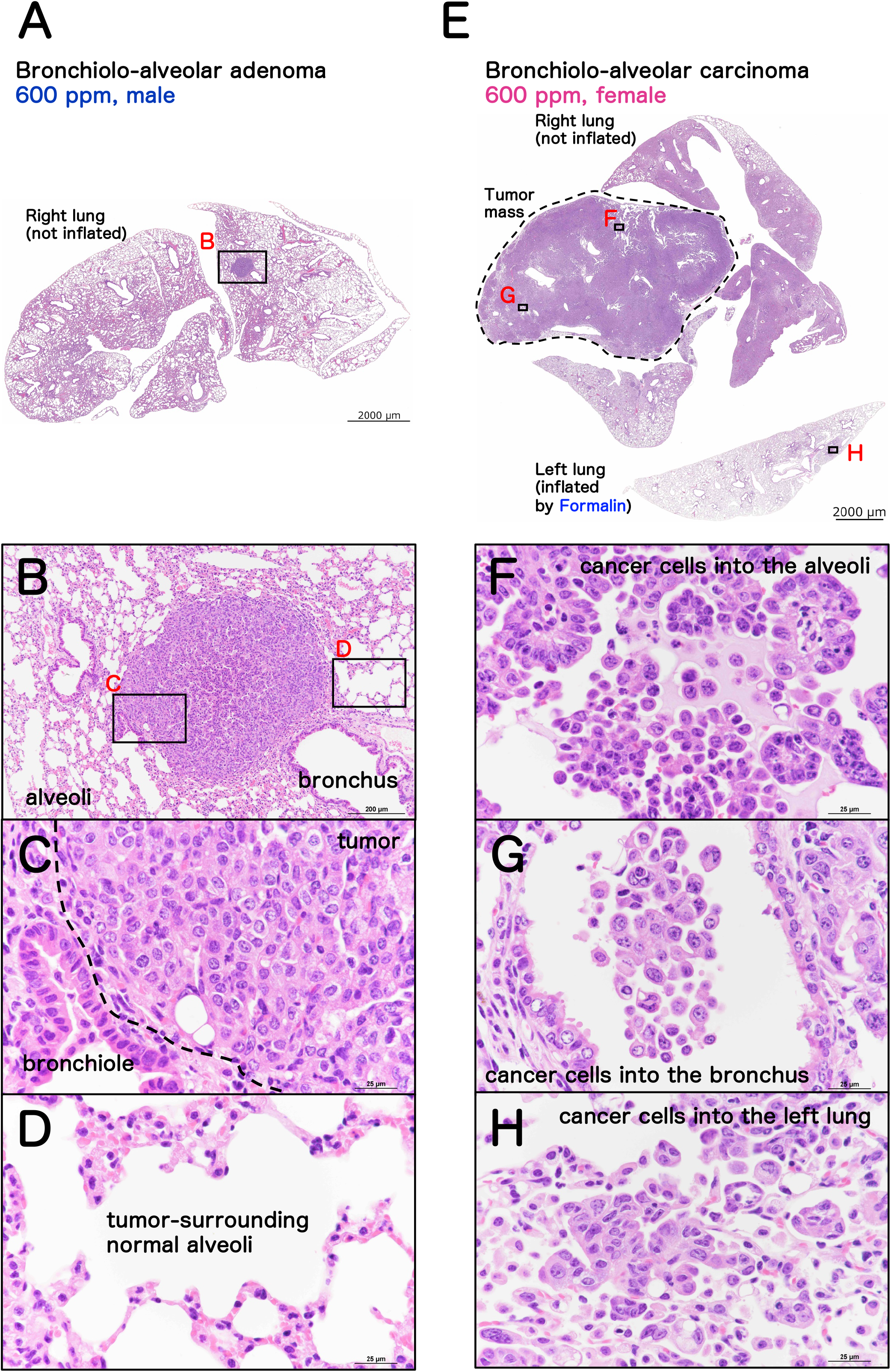
Representative microscopic photographs of the male rasH2 mouse lung tumors. Bronchiolo-alveolar adenoma (A, B, C, D) and carcinoma (E, F, G, H).

**Table 1.**
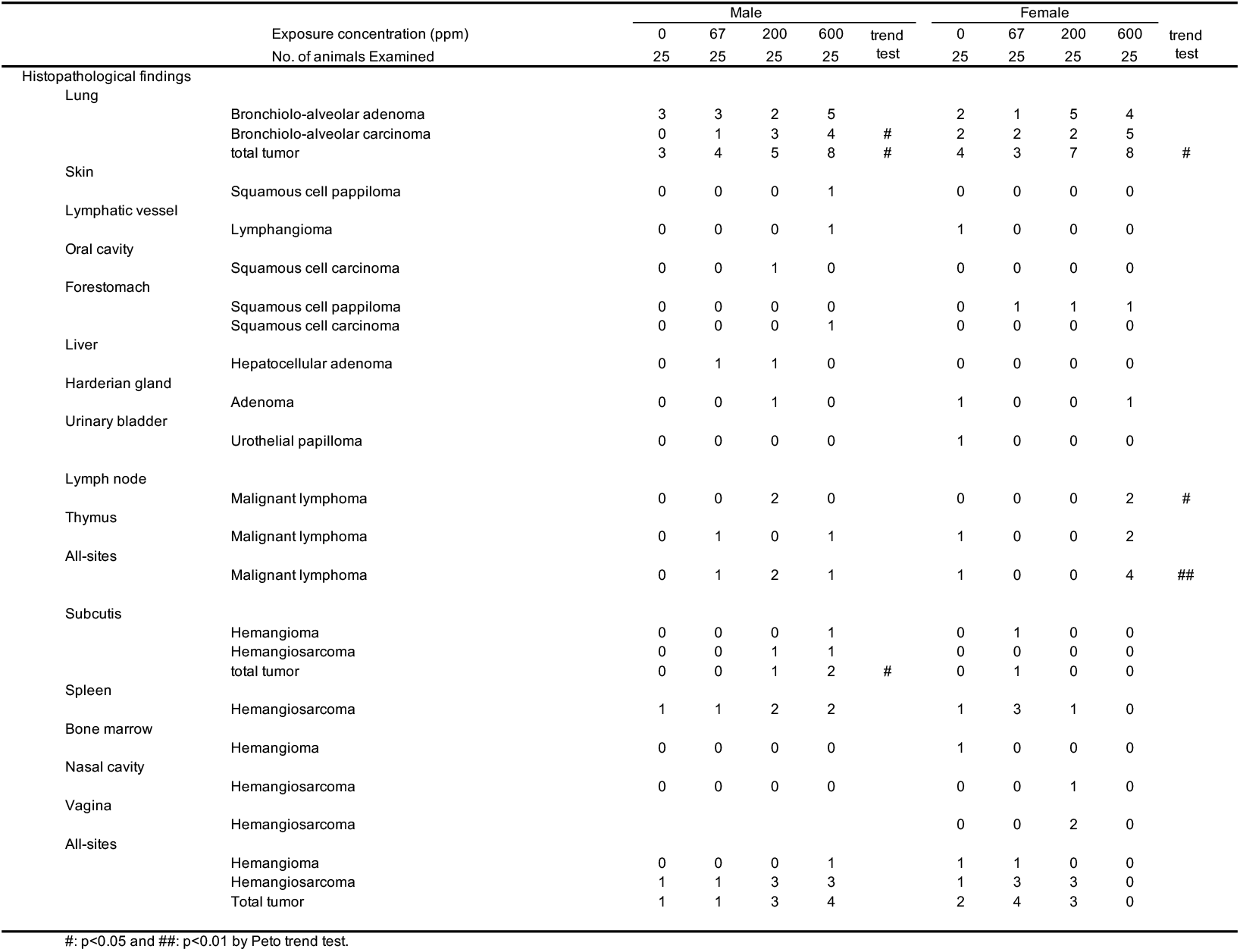
Incidence of the histopathological findings of neoplastic lesions in the all organs.

|  |  | Male |  |  |  |  | Female |  |  |  |  |
| --- | --- | --- | --- | --- | --- | --- | --- | --- | --- | --- | --- |
| Exposure concentration (ppm) |  | 0 | 67 | 200 | 600 | trend | 0 | 67 | 200 | 600 | trend |
| No. of animals Examined |  | 25 | 25 | 25 | 25 | test | 25 | 25 | 25 | 25 | test |
| Histopathological findings |  |  |  |  |  |  |  |  |  |  |  |
| Lung |  |  |  |  |  |  |  |  |  |  |  |
|  | Bronchiolo-alveolar adenoma | 3 | 3 | 2 | 5 |  | 2 | 1 | 5 | 4 |  |
|  | Bronchiolo-alveolar carcinoma | 0 | 1 | 3 | 4 | # | 2 | 2 | 2 | 5 |  |
|  | total tumor | 3 | 4 | 5 | 8 | # | 4 | 3 | 7 | 8 | # |
| Skin |  |  |  |  |  |  |  |  |  |  |  |
|  | Squamous cell papilloma | 0 | 0 | 0 | 1 |  | 0 | 0 | 0 | 0 |  |
| Lymphatic vessel |  |  |  |  |  |  |  |  |  |  |  |
|  | Lymphangioma | 0 | 0 | 0 | 1 |  | 1 | 0 | 0 | 0 |  |
| Oral cavity |  |  |  |  |  |  |  |  |  |  |  |
|  | Squamous cell carcinoma | 0 | 0 | 1 | 0 |  | 0 | 0 | 0 | 0 |  |
| Forestomach |  |  |  |  |  |  |  |  |  |  |  |
|  | Squamous cell papilloma | 0 | 0 | 0 | 0 |  | 0 | 1 | 1 | 1 |  |
|  | Squamous cell carcinoma | 0 | 0 | 0 | 1 |  | 0 | 0 | 0 | 0 |  |
| Liver |  |  |  |  |  |  |  |  |  |  |  |
|  | Hepatocellular adenoma | 0 | 1 | 1 | 0 |  | 0 | 0 | 0 | 0 |  |
| Harderian gland |  |  |  |  |  |  |  |  |  |  |  |
|  | Adenoma | 0 | 0 | 1 | 0 |  | 1 | 0 | 0 | 1 |  |
| Urinary bladder |  |  |  |  |  |  |  |  |  |  |  |
|  | Urothelial papilloma | 0 | 0 | 0 | 0 |  | 1 | 0 | 0 | 0 |  |
| Lymph node |  |  |  |  |  |  |  |  |  |  |  |
|  | Malignant lymphoma | 0 | 0 | 2 | 0 |  | 0 | 0 | 0 | 2 | # |
| Thymus |  |  |  |  |  |  |  |  |  |  |  |
|  | Malignant lymphoma | 0 | 1 | 0 | 1 |  | 1 | 0 | 0 | 2 |  |
| All-sites |  |  |  |  |  |  |  |  |  |  |  |
|  | Malignant lymphoma | 0 | 1 | 2 | 1 |  | 1 | 0 | 0 | 4 | ## |
| Subcutis |  |  |  |  |  |  |  |  |  |  |  |
|  | Hemangioma | 0 | 0 | 0 | 1 |  | 0 | 1 | 0 | 0 |  |
|  | Hemangiosarcoma | 0 | 0 | 1 | 1 |  | 0 | 0 | 0 | 0 |  |
|  | total tumor | 0 | 0 | 1 | 2 | # | 0 | 1 | 0 | 0 |  |
| Spleen |  |  |  |  |  |  |  |  |  |  |  |
|  | Hemangiosarcoma | 1 | 1 | 2 | 2 |  | 1 | 3 | 1 | 0 |  |
| Bone marrow |  |  |  |  |  |  |  |  |  |  |  |
|  | Hemangioma | 0 | 0 | 0 | 0 |  | 1 | 0 | 0 | 0 |  |
| Nasal cavity |  |  |  |  |  |  |  |  |  |  |  |
|  | Hemangiosarcoma | 0 | 0 | 0 | 0 |  | 0 | 0 | 1 | 0 |  |
| Vagina |  |  |  |  |  |  |  |  |  |  |  |
|  | Hemangiosarcoma |  |  |  |  |  | 0 | 0 | 2 | 0 |  |
| All-sites |  |  |  |  |  |  |  |  |  |  |  |
|  | Hemangioma | 0 | 0 | 0 | 1 |  | 1 | 1 | 0 | 0 |  |
|  | Hemangiosarcoma | 1 | 1 | 3 | 3 |  | 1 | 3 | 3 | 0 |  |
|  | Total tumor | 1 | 1 | 3 | 4 |  | 2 | 4 | 3 | 0 |  |

#: p<0.05 and ##: p<0.01 by Peto trend test.

There was a statistically significant increase in the development of bronchiolo-alveolar carcinomas and total lung tumors in male mice exposed to 2-BP and in the development of total lung tumors in female mice exposed to 2-BP (Peto’s trend test).

In addition, there was a statistically significant increase in the occurrence of total hemangiogenic tumors in the subcutaneous tissue of males and malignant lymphomas in the lymph node and all-sites of females (Peto’s trend test). As with the lung tumors, there were no differences in the histological characteristics of these tumors due to 2-BP exposure.

These results indicate that 2-BP is carcinogenic in male and female rasH2 mice exposed to 2-BP by inhalation.

### Reproductive toxicity of 2-BP in rasH2 mice

Previous reports have shown that 2-BP is toxic to the testis ^10^ and ovary ^14^ and to fetal development ^33^ in rodents. Therefore, we examined the toxicity of 2-BP to reproductive organs in male and female rasH2 mice. At necropsy, small testes were observed in 1 male exposed to 200 ppm and in 24 males exposed to 600 ppm, but not in the control males. A summary of the effect of 2-BP on the testes and ovaries is shown in Fig. 5, and absolute and relative weights of various organs are shown in Tables S1 and S2. In male testes, there was a marked and statistically significant concentration-dependent reduction in the testicular weight of 2-BP exposed mice (Fig. 5A, B). Histopathologically, there was a decrease/loss of germ cells and spermatozoa in the seminiferous tubules of the testes and an increase in Sertoli cells and Leydig cells (Fig. 5E). Furthermore, consistent with the histopathological changes in the testis, in the epididymis, a decrease/loss of spermatozoa and an increase in cell debris were observed in the head and tail regions (Fig. 5F). In particular, seminiferous tubular atrophy and Leydig cell proliferation in the testis and reduced sperm with debris in the epididymis were observed in all males exposed to 600 ppm. Grading of these histopathological findings showed a statistically significant increase in testicular and epididymal lesions (Fig. 5G, H). These results indicate that exposure to 2-BP has severe testicular toxicity in male rasH2 mice.

**Fig. 5.**
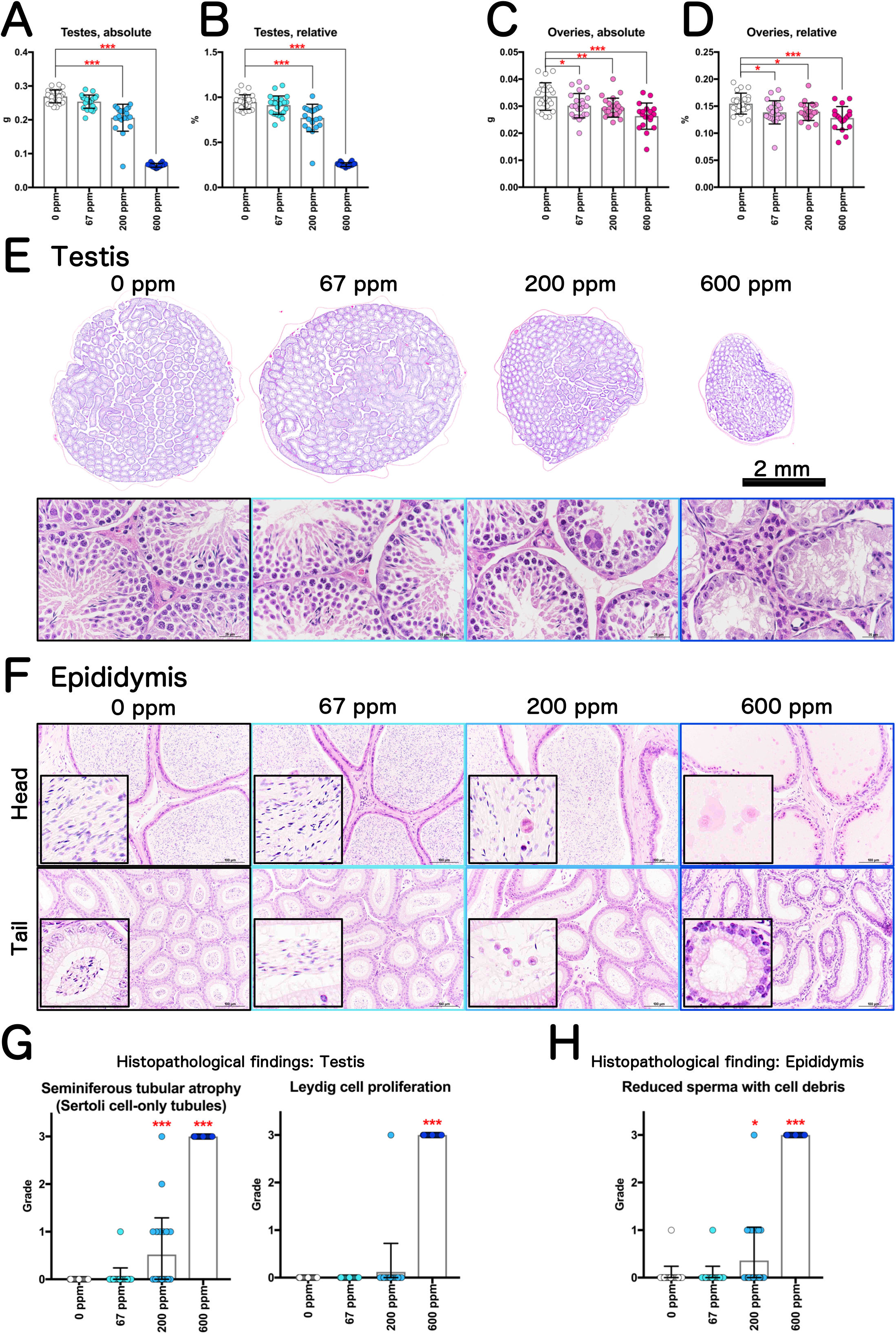
Representative reproductive organ toxicity of rasH2 mice following inhalation exposure to 2-BP (67, 200, or 600 ppm, 6 hours/day, 5 days/week, 26 weeks). Testis organ weights (A, B) and ovary organ weights (C, D) are shown. Loupe images of testes exposed to 2-BP and magnified images (E). Typical histology of the head and tail of the epididymis (F). Summary of the results of various pathological findings of the testes (G) and epididymis (H) observed after 2-BP exposure. Dunnett’s multiple comparison test was used to compare the testes and ovary weights with age-matched control (0 ppm) groups: \*\**p*<0.01 and \*\*\**p*<0.001. Significant difference: *, p<0.05; **, p<0.01; ***, p<0.001 by Chi square test compared with the respective controls for histological grading.

A statistically significant decrease in ovarian weight was observed in female ovaries in a 2-BP exposure concentration-dependent manner (Fig. 5C, D). However, the decrease was mild and no histopathological changes were observed. Therefore, at the concentration used in this study, 2-BP had little or no ovarian toxicity in rasH2 female mice.

### Hematopoietic toxicity of 2-BP in rasH2 mice

Previous reports have shown that 2-BP has hematopoietic toxicity in male and female rodents. Therefore, we also examined the toxicity of 2-BP to hematopoietic organs in male and female rasH2 mice. Figure 6A shows representative photographs of the bone marrow in control and 600 ppm exposed mice, spleen weights are shown in Fig. 6B-E, and hematologic and blood biochemistry data are shown in Tables S3 and S4. Significant decreases in platelet counts were commonly observed in the male and female groups exposed to more than 200 ppm and significant decreases in erythrocyte counts and high MCH and MCV levels were commonly observed in the male and female 600 ppm groups. However, histopathological observations of bone marrow showed no change in 2-BP exposed male or female rasH2 mice (Fig. 6A). A significant decrease in absolute spleen weight was observed only in the female 600 ppm exposure group, but no change was observed in relative weight of the spleen in females, and no change in spleen weight was observed in males (Fig. 6B-E). Histopathological observations of the spleen showed no significant changes due to 2-BP exposure (Fig. 6F). In the bone marrow and spleen, pathological findings related to hematopoietic cells showed no effect of 2-BP exposure (Fig. 6G, H). These results indicate that 2-BP did not exhibit hematopoietic toxicity in male or female rasH2 mice in this study.

**Fig. 6.**
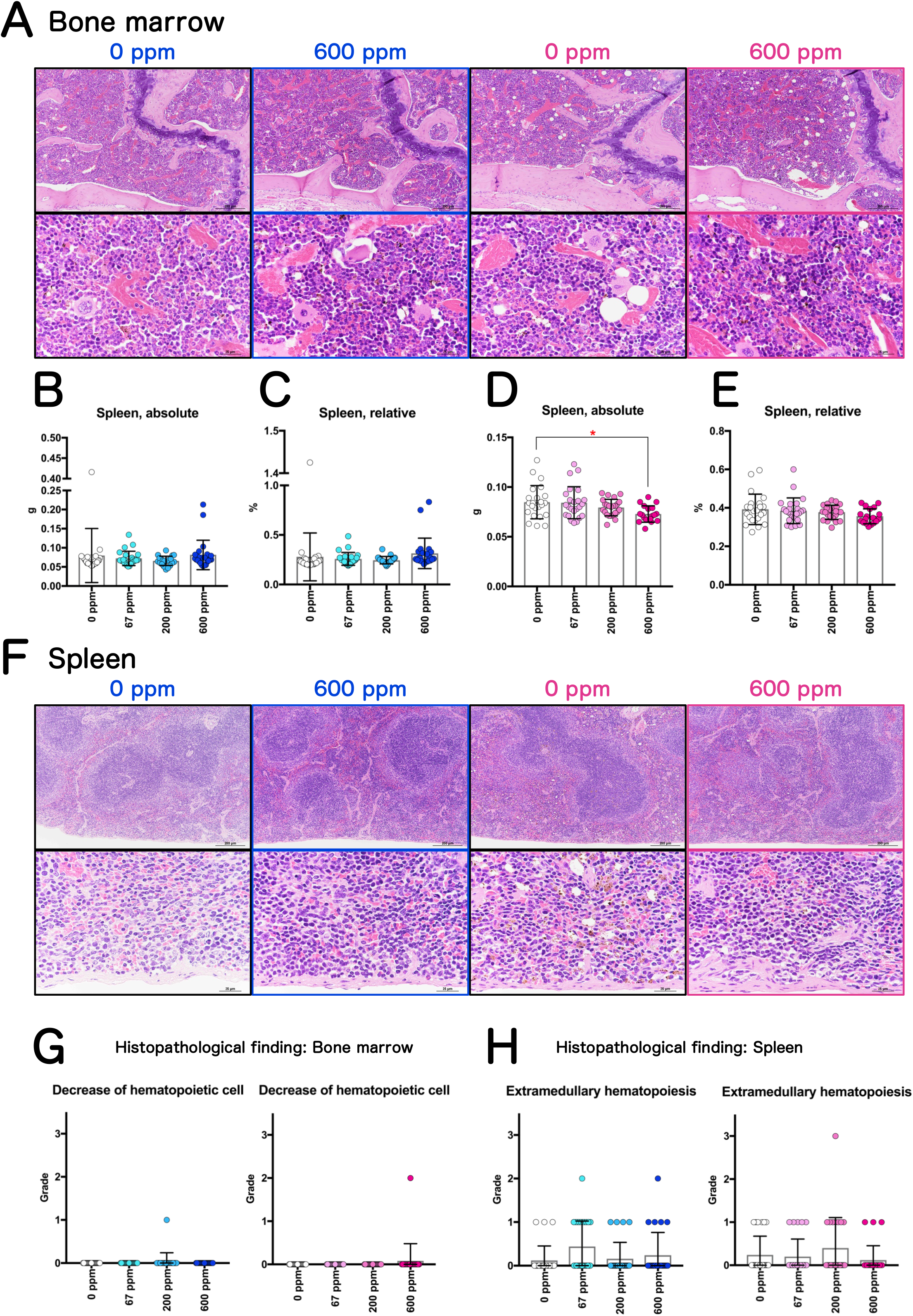
Representative hematopoietic organs toxicity of rasH2 mice following inhalation exposure to 2-BP (67, 200, or 600 ppm, 6 hours/day, 5 days/week, 26 weeks). Typical histology of the bone marrow (A). Spleen weights of males (B, C) and females (D, E). Typical histology of the spleen (F). Summary of the results of various pathological findings of the bone marrow (G) and spleen (H) observed after 2-BP exposure. Dunnett’s multiple comparison test was used to compare the spleen weights with the age-matched control (0 ppm) groups: \**p*<0.05.

## Discussion

In this study, we investigated various toxicological effects of systemic inhalation exposure of up to 600 ppm 2-Bromopropane (2-BP) by male and female rasH2 mice. The results showed a concentration-dependent increase in lung tumor development in both male and female rasH2 mice, which was significant by Peto’s trend test. Furthermore, 2-BP was found to have testicular toxicity. This study provides the first evidence that inhalation of 2-BP is carcinogenic in male and female mice and is a testicular toxin in the rasH2 mouse model (Fig. 7).

**Fig. 7.**
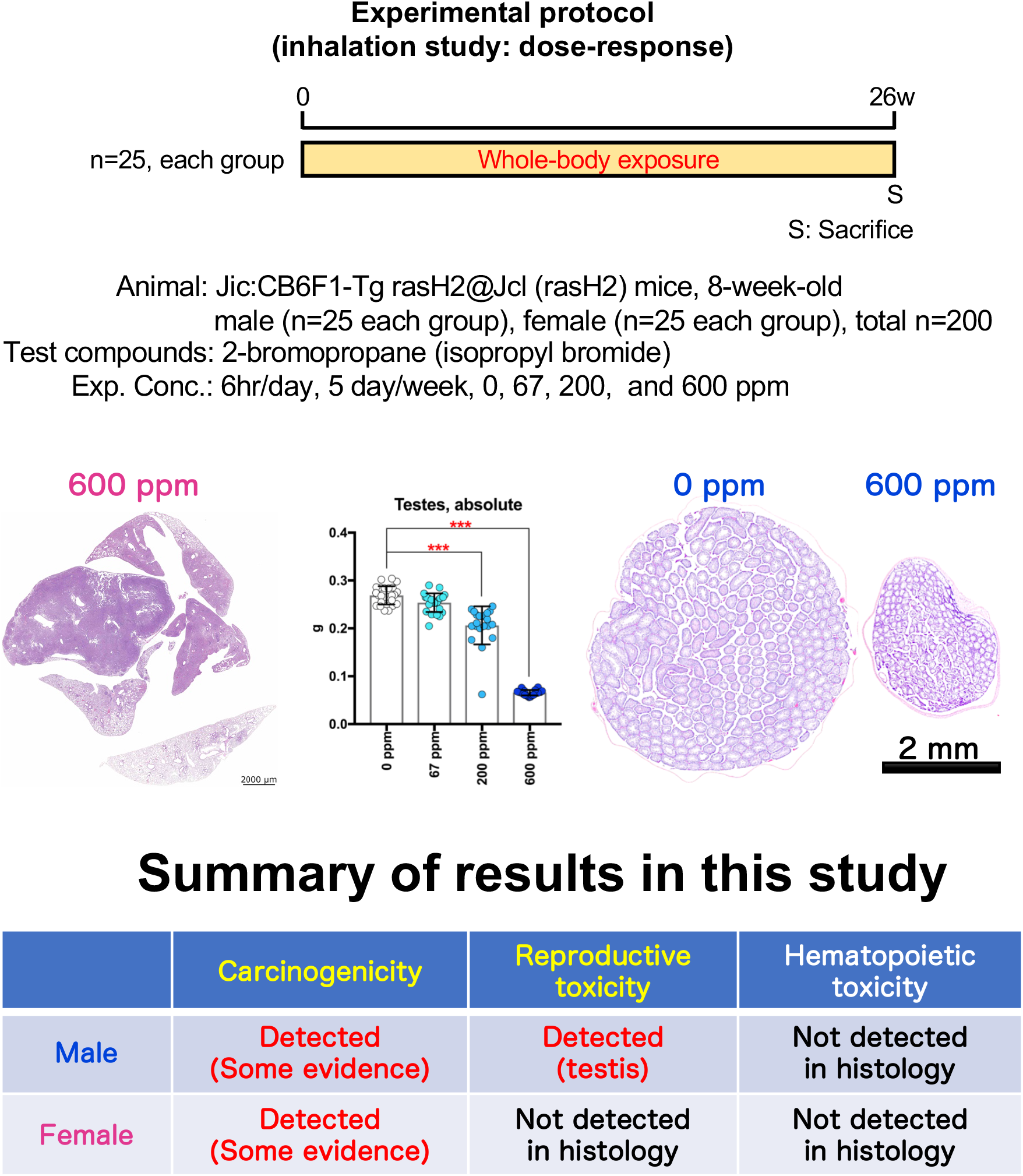
Graphical abstract of this study.

The reported incidence of total lung tumors (adenomas plus carcinomas) in the lungs of control rasH2 mice from the breeding colony established in 2018 at CLEA Japan is approximately 7 - 20%^34^. In the present study, the total lung tumor incidence in the control groups were 3/25 (12%) in males and 4/25 (16%) in females, which is comparable to the background total lung tumor incidence reported by CLEA Japan. In addition, the histopathological features of the lung tumors observed in this study were not qualitatively altered by 2-BP exposure. Therefore, we can conclude that the increase in tumorigenesis in the lungs of rasH2 mice was caused by inhalation exposure to 2-BP.

The mechanism of 2-BP carcinogenesis remains to be investigated in future studies. 1-Bromopropane (1-BP), an alkane bromide with a chemical structural formula similar to that of 2-BP, is also carcinogenic and was classified as Group 2B (possibly carcinogenic to humans) by IARC in 2016 ^35,36,37^. While the carcinogenic mechanism of 1-BP has not been fully elucidated, it has been considered to be a nongenotoxic carcinogen based on negative *in vitro* and *in vivo* mutagenicity studies ^36,38^. In the IARC Monograph 115, evaluation of the mechanisms of carcinogenesis is based on the “key characteristics of carcinogens” proposed by Smith *et al*. ^39^. There is strong evidence that 1-BP is electrophilic or can be metabolically activated and induces oxidative stress, chronic inflammation, and is immunosuppressive ^36^. It is possible that the mechanisms of carcinogenesis of 2-BP may have similarities to those of 1-BP, however, this remains to be determined in future studies.

As noted above, 1-BP is considered to be a nongenotoxic carcinogen. Maeng et al., 1997,^19^ reported that 2-BP was mutagenic in Salmonella typhimurium TA100 and TA1535, but that mutagenicity using Escherichia coli WP2 uvrA and chromosomal aberration tests using Chinese hamster lung cells were negative. Furthermore, micronuclei were not increased in experiments in which 2-BP was administered intraperitoneally to F344 rats at doses up to 500 mg/kg BW once daily for 28 days^19^. Therefore, the mutagenicity of 2-BP remains to be determined. In order to confirm mutagenicity as a carcinogenic mechanism of 2-BP, the results of *in vivo* mutagenicity assays using gpt delta mice, and using organs in which a cancer develops are needed ^40–42^.

Based on previous reports, 2-BP has been shown to have numerous testicular toxic effects in humans and rats. Decreased sperm counts and decreased active sperm rates have been observed in male workers exposed to a 2-BP cleaning solution at an electronic factory in South Korea ^2,3^. A report on the effect of exposure to low concentrations of 2-BP at a 2-BP manufacturing plant also suggested the possibility that exposure to 2-BP resulted in decreased sperm count and motility ^43^. Moreover, testicular toxicity has been observed in various experiments in which rats were exposed to 2-BP at concentrations up to 3000 ppm by inhalation ^5,44^, intraperitoneal administration of up to 500 mg/kgBW ^7^, oral administration of up to 3.5 g/kgBW/day ^10^, and subcutaneous administration of up to 1800 mg/kgBW ^45^. It has been reported that the target cells of 2-BP in the testis are spermatogonia, and that apoptosis is induced via Bcl2 family genes and the Fas signaling system ^11,12,46,47^. Consistent with these results, in the present study, dramatic testicular atrophy was observed in male mice exposed to 600 ppm 2-BP, indicating that 2-BP is a potent testicular toxicant in rasH2 mice. In contrast to the effects on the testes, exposure to 2-BP resulted in a significant, but mild, concentration-dependent decrease in ovarian weight, but no histopathological changes were observed in the ovaries. Therefore, it was determined that no histological ovarian toxicity was observed in rasH2 female mice in the present study at any of the exposure concentrations.

In workers exposed to 2-BP, hematopoietic toxicity with anemia as the symptom has been reported ^2,3,43^. Based on these reports, we evaluated the hematopoietic toxicity of 2-BP exposure in rasH2 mice. Analysis of peripheral blood samples at autopsy revealed that a significant decrease in erythrocyte count was observed in the 600 ppm groups of male and female rasH2 mice. However, no anemia symptoms were observed, and histopathological analysis of bone marrow and spleen showed no effects of 2-BP exposure. Based on these results, it was concluded that 2-BP did not exhibit hematopoietic toxicity histologically in the present study.

In the present study we identified 2-BP as a potential carcinogen. However, the use of the rasH2 mouse model did not allow us to elucidate the details of the target organs or the mechanisms of carcinogenesis in the mouse. In addition, a 2-year carcinogenicity study in rats, another rodent species, is needed to ascertain the carcinogenicity of 2-BP in test animals and potentially in humans. We are currently in the process of compiling the results of a carcinogenicity study of systemic inhalation exposure to 2-BP in F344 rats.

## Conclusions

In this study, various toxic effects of 2-Bromopropane (2-BP), including carcinogenicity, were investigated in male and female rasH2 mice exposed by whole-body inhalation to 67, 200 and 600 ppm. The results showed a concentration-dependent increase in lung tumor development in male and female rasH2 mice, and this increase was significant by Peto’s trend test. Furthermore, in male rasH2 mice, 2-BP was found to be testicular toxin. This study is the first to demonstrate that 2-BP is carcinogenic in male and female mice and is a testicular toxin in male mice in the rasH2 mouse model. Therefore, our results identify 2-BP as a potential carcinogen and demonstrate that additional carcinogenic studies on 2-BP need to be carried out. This will provide important information for the IARC working group in evaluating the carcinogenicity of 2-BP.

## Abbreviations

1-BP: 1-Bromopropane
2-BP: 2-Bromopropane
CFC: Chlorofluorocarbon
HE: hematoxylin and eosin
IARC: International Agency for Research on Cancer
OECD: Organisation for Economic Co-operation and Development
rasH2: Jic:CB6F1-Tg ras H2@Jcl

## Data availability

The datasets used during the current study are available from the corresponding author on reasonable request.

## Acknowledgments

We wish to thank Dr. David B. Alexander of Nanotoxicology project, Nagoya City University Graduate School of Medicine for his insightful comments and English editing. Finally, we would like to express our heartfelt gratitude to all the Japan Bioassay Research Center staff.

## Author information

### Contributions

A.S. performed the experiments and analyzed the data. M. S., K. T., H. S. and Y. U. assisted with animal experiment including exposure animal care and sacrifices. K.T., H.S., Y.U. and S.Y. performed histopathological diagnoses. Y.G., S.Y. and Y.U. drafted and revised the manuscript. All authors approved the manuscript as submitted.

### Corresponding author

Shotaro Yamano

